# The chromatin accessibility dynamics during cell fate specifications in zebrafish early embryogenesis

**DOI:** 10.1101/2023.10.13.562312

**Authors:** Qiushi Xu, Yunlong Zhang, Wei Xu, Dong Liu, Wenfei Jin, Xi Chen, Ni Hong

**Affiliations:** Harbin Institute of Technology, Harbin, China; Shenzhen Key Laboratory of Gene Regulation and Systems Biology, Department of Systems Biology, School of Life Sciences, Southern University of Science and Technology, Shenzhen 518055, Guangdong, China; GMU-GIBH Joint School of Life Sciences, The Guangdong-Hong Kong-Macau Joint Laboratory for Cell Fate Regulation and Diseases, Guangzhou Medical University, Guangdong, China

## Abstract

Chromatin accessibility plays a critical role in the regulation of cell fate decisions. Although gene expression changes have been extensively profiled at the single-cell level during early embryogenesis, the dynamics of chromatin accessibility at *cis*-regulatory elements remain poorly studied. Here, we used a plate-based single-cell ATAC-seq method to profile the chromatin accessibility dynamics of over 10, 000 nuclei from zebrafish embryos. We investigated several important time points immediately after zygotic genome activation (ZGA), covering key developmental stages up to dome. The results revealed key chromatin signatures in the first cell fate specifications when cells start to differentiate into enveloping layer (EVL) and yolk syncytial layer (YSL) cells. Finally, we uncovered many potential cell-type specific enhancers and transcription factor motifs that are important for the cell fate specifications.

## Introduction

In early embryogenesis, the totipotent zygote first enters the cleavage stage during which a series of synchronised mitotic divisions occur (1). The embryo is transcriptionally quiescent, and the progress is mainly driven by maternal factors. The main wave of zygotic genome activation (ZGA) when the genome of the embryo begins to make their own RNAs and proteins only occurs at later cell cycles (2). Starting from ZGA, embryonic cells become asynchronous along developmental trajectories. They undergo a series of complex events, such as cellular differentiation and migration, and eventually gives rise to different cell types with committed fates and distinct functions (3). Understanding how cells acquire specific fates is one of the key goals in developmental biology. Due to the heterogeneous and asynchronous nature of the developmental processes, single-cell genomics methods are necessary and serve as powerful approaches for the unbiased investigation of molecular profiles of each cell during development.

In zebrafish, recent single-cell RNA-seq (scRNA-seq) developmental atlases provided gene expression overviews of all major cell types from embryonic to adult stages (4–10). For example, the first two zebrafish scRNA-seq atlases (4, 5) described in great details the gene expression patterns during embryogenesis at unprecedented temporal and cellular resolution, revealing that cells exhibit potential plasticity during development and cell fate specification is a gradual and non-binary process. Furthermore, those transcriptomic profiling studies revealed stage-specific gene expression programs, branching points of emerging cell fates and key regulators during different developmental trajectories (4–10). While those datasets serve as invaluable resources to study organismal development, the investigation regarding the regulatory events underlying those transcriptomic and cell fate changes remains limited.

The progressive cell fate specifications are driven by the interaction between transcription factors and *cis*-regulatory elements located in regions such as promoters and enhancers (11). In general, chromatin around *cis*-regulatory elements needs to be in an accessible state to allow transcription factor binding. Therefore, genome-wide investigation of chromatin accessibility will provide insights on the molecular mechanisms of cell fate commitments. In recent years, several studies used the assay for transposase-accessible chromatin using sequencing (ATAC-seq) to interrogate the chromatin accessibility landscapes during zebrafish early embryogenesis (12–14). Those studies generated detailed chromatin dynamics during early development, demonstrating the gradual increase of chromatin accessibility around *cis*-regulatory elements on a global scale.

To date, most chromatin accessibility studies during zebrafish early embryogenesis have been performed at the bulk level (12–14). A limited number of single-cell chromatin accessibility studies mainly focused on the post-gastrula stages or adult tissues (15–17). Therefore, it is still not clear if and when cells become different at the chromatin level during early embryogenesis. In this study, we used a plate-based single-cell ATAC-seq (scATAC-seq) method (18, 19) to investigate the chromatin accessibility states of 10, 518 single embryonic nuclei from ZGA up to the dome stage. Different accessible chromatin landscapes among cells begin to emerge when the embryo is transitioning from the sphere to the dome stage. Distinct chromatin states of the enveloping layer (EVL), the yolk syncytial layer (YSL) and the deep layer (DEL) cells are clearly observed. Compared to DEL cells, the EVL and YSL cells obtain their own signature profiles where specific distal regions that are putative enhancers change from an inaccessible to accessible state. Overall, our time-course single-cell data provides valuable information about molecular processes underlying the first cell fate specification.

## Materials & Methods

### Zebrafish maintenance

The wild-type TU (Tübingen) zebrafish were maintained at 28°C with 14/10 hours of light/dark cycles, respectively. Embryos were collected and transferred to the zebrafish embryo medium (5 mM NaCl, 0.17 mM KCl, 0.33 mM CaCl2, 0.33 mM MgSO4, and 0.002% methylene blue) at 28°C immediately after fertilization. All animal experiments were performed with approval from the Experimental Animal Welfare Ethics Committee, Southern University of Science and Technology.

### Embryo collection and cell dissociation

Embryos at different time points after fertilisation were collected, including 3 hours post fertilisation (hpf) (the 1k-cell stage), 3.3 hpf (the high stage), 3.7 hpf (the oblong-sphere stage), 4.1 hpf (the sphere-dome stage) and 4.3 hpf (the dome stage). Embryos showing inconsistent developmental status were excluded. A total of 50 to 100 embryos in each stage were collected for one batch of experiment. To get embryonic cells, chorions were removed by treating with 1 mg/mL Pronase E (Solarbio, P8360) at room temperature for 3 min. Then the embryos were washed three times with 10 mL 1× PBS and manually deyolked using tweezers under a microscope, and cells were dispersed by gentle pipetting. The dispersed cells were transferred to a 1.5 mL tube and collected by centrifugation at 500g, 4°C for 5 minutes. The pellet was resuspended in 1×PBS-0.5% bovine serum albumin (BSA), and the cell number was counted using a C-Chip disposable hemocytometer (INCYTO, cat. no. DHC-N01) after stained with trypan blue (Sigma-Aldrich, T8154).

### Single-cell ATAC-seq

The scATAC-seq libraries were prepared using a plate-based workflow as described previously (18, 19). Briefly, 50, 000 cells were lysed in 50 μL RSB-DTN (10 mM Tris-HCl, pH 7.4, 10 mM NaCl, 3 mM MgCl2, 0.1% IGEPAL CA-630, 0.1% Tween-20 and 0.01% digitonin) on ice for 3 minutes. The lysate was then washed with 950 μL cold RSB-T (10 mM Tris-HCl, pH 7.4, 10 mM NaCl, 3 mM MgCl2, and 0.1% Tween-20) and centrifuged at 1, 000g, 4°C for 10 minutes. The cell pellet was resuspended in 50 μL tagmentation mix (33 mM Tris-acetate pH 7.8, 66 mM potassium acetate, 10 mM magnesium acetate, 16% dimethylformamide, 0.01% digitonin, and 5 μL Tn5) and incubated at 37 °C for 30 minutes with gentle shaking. The reaction was stopped by adding 50 μL 2× stop buffer (10 mM Tris-HCl pH 8.0, 20 mM EDTA pH 8.0). Nuclei were centrifuged at 500 g, 4 °C for 5 minutes and then resuspended in 500 μL 0.5× PBS-0.5% BSA, stained by adding 1.5 μL 1 μg/μL DAPI and hold on ice. The single DAPI-positive nucleus was sorted into each individual well of a 384-well plate containing 3 μL preloaded lysis buffer (11 mM Tris-HCl pH 8.0, 11 mM NaCl, 0.22% SDS, 10 μM Nextra_S5_short primer, and 10 μM N7xx primer), and incubated at 65 °C for 15 minutes. After a brief centrifugation, 2 μL of 10% Tween-20 and 5 μL NEBNext High-fidelity 2× PCR Master Mix (NEB, M0541) was added to each well sequentially. Pre-amplification in each well was performed as follows: 72°C for 5 minutes; 98 °C for 1 minute; 10 cycles of 98 °C for 10 seconds, 63°C for 30 seconds, 72°C for 20 seconds. All pre-amplified libraries in each well from the same 384-well plate were pooled together and transferred into a 15-mL tube. The pooled pre-amplified libraries were purified by the QIAquick PCR purification kit (Qiagen, 28104). Excessive primers were digested by Thermolabile Exonuclease I (NEB, M0568) at 37 °C for 30 min, and then purified by 1.5× VAHTS DNA clean beads (Vazyme, N411-01) and eluted in 20 μL of nuclease-free water. The pooled pre-amplified libraries from each plate were mixed with 2.5 μL scATAC_S5xx primer (10 μM), 2.5 μL Illumina P7 primer (10 μM), and 25 μL NEBNext High-Fidelity 2× PCR Master Mix, followed by a final PCR for amplification: 98 °C for 1 minute; 12 cycles of 98 °C for 10 seconds, 63 °C for 30 seconds, 72 °C for 20 seconds. The final libraries with plate-specific barcodes were purified by 1.2× VAHTS DNA clean beads and eluted in 20 μL of nuclease-free water. Equimolar libraries from different plates were pooled and sequenced on the Illumina HiSeq X-Ten machine using standard sequencing scheme.

### Data preprocessing

After sequencing, BCL files were converted to FastQ files using bcl2fastq (v.2.19) from Illumina. Read 1 and Read 2 FastQ files were processed using a Snakemake pipeline (20). In brief, 3′-end adapter sequence was trimmed with fastp (21). Then the trimmed reads were mapped to UCSC danRer11 reference genome using hisat2 (22). Reads with mapping quality less than 30 were removed by samtools (23). Duplicated reads were removed using the MarkDuplicates function of the Picard tool (http://broadinstitute.github.io/picard/). Reads from single cells were merged together and deduplicated again using samtools and Picard to generate a merged BAM file for each stage. Peak calling of the merged BAM files was performed using MACS2 (24). The peak-by-cell matrix was generated for each stage by counting the number of reads within peaks from individual cells with bedtools (25). The software and packages used in this pipeline can be found in the “Snakefile” provided in the GitHub repository(https://github.com/dbrg77/scATAC_snakemake).

### Quality control (QC)

To remove low-quality cells, we computed four quality control (QC) metrics of each cell: (1) The number of unique nuclear fragments: the number of reads mapped to the nuclear genome after deduplication; (2) Mapping rate: the overall alignment rate calculated by hisat2; (3) Nucleosome signal: the ratio of mono-nucleosomal fragments (147 - 294 bp) to nucleosome-free fragments (<147 bp) which was calculated by the “NucleosomeSignal” function from the R package Signac (26); (4) Transcriptional start site (TSS) enrichment score: a ratio between aggregated distribution of reads centred on the TSS and the flanking regions, which was calculated by the “TSSErichment” function of Signac. Based on the per-stage distribution of each QC metric, the following thresholds were used for different stages to ensure that cells in earlier stages with lower accessibility would not be removed: 3 hpf (unique nuclear fragments > 1000 & mapping rate > 70 & TSS enrichment > 1), 3.3 hpf and 3.7hpf (unique nuclear fragments > 2000 & mapping rate > 70 & TSS enrichment > 1.5), 4.1 hpf (unique nuclear fragments > 2000 & mapping rate > 75 & TSS enrichment > 1.5), 4.3 hpf (unique nuclear fragments > 2000 & mapping rate > 75 & TSS enrichment > 2).

### Normalization and 2D visualisation

We used the “reduce” function of the R package GenomicRanges (27) to combine all intersecting peaks between stages and created a unified set of peaks. To generate a genomic blacklist, we used MACS2 to perform peak calling on a prior “input” dataset of ATAC-seq experiment where Tn5 was used to cut zebrafish naked genomic DNA (14). Peaks that intersected with the blacklist were removed. Based on the remaining peaks, we created a peak-by-cell count matrix. The count matrix was normalized them using the term frequency–inverse document frequency (TF–IDF) transformation. Then singular value decomposition (SVD) was applied to the matrix to obtain a lower-dimensional representation. Subsequently, the 2 - 30 latent semantic indexing (LSI) components were used to perform UMAP to visualise the final result in 2-dimension plots.

### Data integration of different stages

To integrate data from all stages, we used BBKNN (28) for batch correlation. We created the combined peak-by-cell matrix with all stages and performed the LSI analysis using the same workflow described above. The top 20 LSI components were used as an input to the bbknn function from the Python package Scanpy (29). To integrate data for later stages (4.1 hpf and 4.3 hpf) for a detailed analysis of cell types, we used the “IntegrateEmbeddings” function of Signac to integrate data from 4.1 hpf (sphere-dome) and 4.3 hpf (dome) stages and projected them into a shared low-dimensional space. After that, new LSI-coordinates (rLSI) of cells were generated, which represented the low-dimensional cell embeddings after integration. We clustered sphere-dome- and dome-stage cells and visualised them using the “FindCluster” (resolution = 0.6) and RunUMAP functions from Signac.

### Identification of peak modules

We used the non-negative matrix factorization (NMF) algorithm from Python scikit-learn toolkit (30) (version 1.1.0) to factorise the normalized peak-cell matrix into two matrices: a matrix H (Peaks by Peak modules), and a matrix W (Peak modules by Cells). The H matrix essentially groups peaks into peak modules such that peaks within the same peak module have similar patterns across single cells, that is, they co-vary among different cells. The number of peak modules is often much lower than the number of original peaks. This is reminiscent of the gene expression programs after performing NMF on a gene expression matrix, where genes that co-vary among cells are grouped into programs. The number of peak modules (*k*) was tested from 3 to 60, and the optimal number was determined to be 25. Modules with high usages in only one batch were possibly caused by the batch effect and ignored for the subsequent analyses. Monotonic increasing (modules 2, 3, 4 and 6) and decreasing module (module 10) during embryo development were identified based on the usage distribution.

### Identification of differentially accessible sites

To identify differentially accessible sites between cell clusters, we performed peak calling of each cell cluster in the 4.1 hpf (sphere-dome) and 4.3 hpf (dome) stages. After combining the peaks and normalising the resulting matrix, we performed differential accessibility (DA) test to determine the differential accessible sites and marker peaks of each cell type by the “FindMakers” function of Seurat (min.pct = 0.25, logfc.threshold = 0.5). Logistic regression (LR) method was chosen in this step. The total number of fragments of each cell were used as a latent variable to mitigate the effect of differential sequence depth on the result.

### Motif analysis

For the *de novo* motif discovery, the 2, 000 peaks from each peak module or EVL or YSL marker peak were used as the input for the “findMotifsGenome.pl” function from HOMER (31) with the parameter “-size 500” to look for the centre 500 bp of each peak. To look at motif deviations across different cells, position frequency matrix (PFM) of non-redundant vertebrates’ transcription factors (TF) binding profiles form the JASPAR database 2022 (32) version was used as the input to the “AddMotifs” function of Signac. Then chromVAR (33) was invoked by the “RunChromVAR” function was used to compute a per-cell motif activity score and created a motif-cell matrix. The differential activity was computed using the “FindMarkers” function.

### Gene activity score calculation

To quantify the gene activities of homeodomain transcription factors, the function “GeneActivity” from Signac was used to generate a gene activity matrix of all homeodomain transcription factors. In this process, gene coordinates were extracted and extended to include the 2 kb region upstream of the transcriptional start site of the gene. Then the number of fragments mapped to the extended region consisting of the 2 kb proximal promoter and the gene body, were counted and treated as the gene activity score for the gene. Finally, the function “AverageExpression” was used to get the average values of the fragment counts for each cell type.

### DNA methylation analysis

We used the publicly available WGBS data set (ArrayExpress, accession number E-MTAB-12535) at the 1k stage to investigate the DNA methylation profiles around LTR regions from module 10 peaks. BigWig files in danRer10 containing DNA methylation values were downloaded. The UCSC genome browser utility “liftOver” was used to convert the coordinates of LTR regions from danRer11 to danRer10. Then the function “computeMatrix” from deepTools (34) were used to calculate methylation signals for given regions with the following option: --skipZeros -a 1000 -b 1000 --binSize 50.

### Co-accessibility analysis

The co-accessibility of peaks was estimated by Cicero (35). We calculated the co-accessibility score for each peak pair using the “run_cicero” function. The peak pairs with a co-accessibility score greater than 0.1 were retained. Next, we generated a co-accessibility network using the “generate_ccans” function based on the co-accessibility score framework. The result was visualized using the “LinkPlot” function from Signac.

### Trajectory analysis

We used a diffusion map (36) to perform trajectory analysis of the 4.1 hpf (sphere-dome) and 4.3 hpf (dome) stage cells. Scores for the top 12 rLSI components were used as the input to compute the neighbourhood graph of cells. Then we generated a diffusion map embedding using the “diffmap” function from Scanpy. To calculate the pseudotime value of each cell, we set DEL1 as the root due to its lower accessibility, and performed diffusion pseudotime (DPT) analysis using the “dpt” function. Finally, we fitted a vector generalized additive model (VGAM) with a degree of freedom of 3 to show the dynamics of peak accessibility in pseudotime using the R package VGAM (37).

## Results

### Single-cell ATAC-seq reveal the accessible chromatin landscape from ZGA to dome in zebrafish

To capture the earliest time point when embryonic cells become different from each other, a plate-based scATAC-seq method previously developed in our labs (18, 19) was used to profile the chromatin accessibility of each single nucleus in the zebrafish embryos (**Figure 1A** and **Supplementary Figure 1A**). Several key developmental time points immediately after ZGA were covered, spanning the 1k, high, oblong, sphere and dome stages. A total of 11, 520 nuclei from staged embryos were profiled and over 3 billion read pairs were obtained with an overall mapping rate of >75% to the zebrafish genome (**Supplementary Figure 1B**). After quality control (see **Methods**; **Supplementary Figure 1C**), 10, 518 nuclei were used for further analysis (**Supplementary Table 1**). After duplication removal, the median numbers of unique fragments mapped to the nuclear genome range from ∼26, 000 to ∼60, 000 (**Supplementary Figure 1B**), which is at the higher end compared to the scATAC-seq atlases in other stages and adult tissues (15–17). Peak calling using MACS2 from merged single nuclei (see **Methods**) identified 446, 501 open chromatin regions. Visual inspections on a genome browser of the read pile-up from aggregated single nuclei within the same time point indicated the data was of good quality (**Figure 1B** and **Supplementary Figure 1D**). The aggregated single-nucleus profiles were also comparable to a recent high-quality bulk ATAC-seq data in the corresponding stages (14).

**Figure 1.**
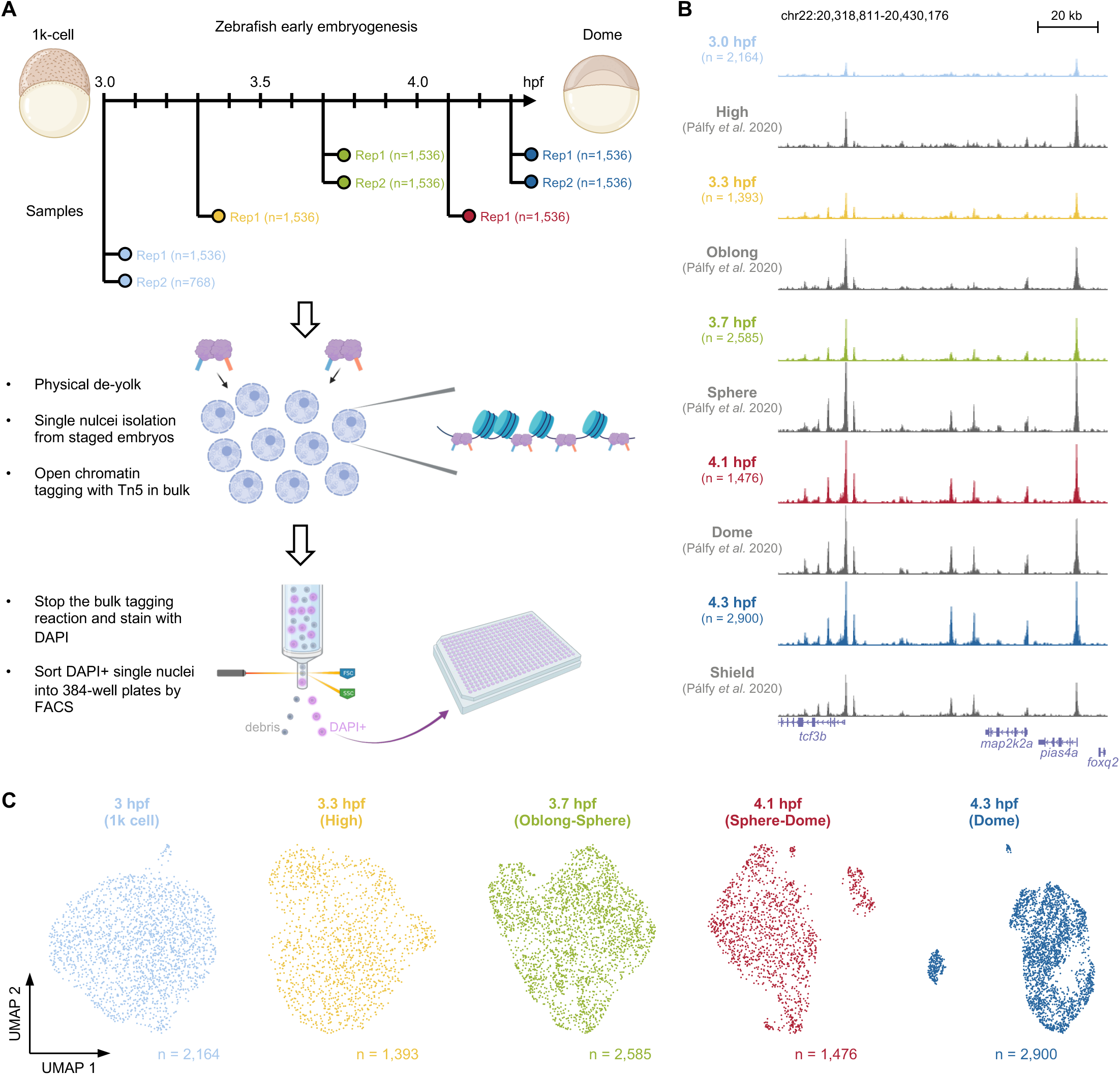
Single-nucleus chromatin accessibility maps of zebrafish early embryogenesis. **(A)** Schematic view of the overall experimental workflow. **(B)** UCSC genome browser tracks of a genomic locus for all time points. The aggregated single-nucleus ATAC-seq profiles per time point was shown together with a previously published bulk ATAC-seq data around similar stages. **(C)** 2D UMAP visualisation of the single-nucleus ATAC-seq data per time point.

Following dimensionality reduction by the latent semantic indexing analysis(26, 38), uniform manifold approximation and projection (UMAP) (39) was used to project the data onto a two-dimensional space for visualisation per stage (**Figure 1C**). No apparent separation of nuclei was observed from 3 to 3.7 hours post fertilisation (hpf), indicating cells were mostly homogeneous at the chromatin level during the first few stages following ZGA. Notably, discrete nuclei clusters started to emerge from 4.1 hpf and onwards (**Figure 1C**). The observation suggested that cells began to acquire distinct accessible chromatin profiles for the first time at the transition from the sphere to the dome stage.

### Key peak modules have different genomic features and dynamics during development

To get an overview of the chromatin accessibility landscape during this developmental period, a batch balanced k-nearest neighbour (BBKNN) method (28) was used to remove batch effect and integrate all stages into a single UMAP space (**Figure 2A** and **Supplementary Figure 2A**). Next, non-negative matrix factorisation (NMF) (40, 41) was utilised to find peak modules that covary across single nuclei throughout the development (**Figure 2B**). The first matrix from the NMF represents how much each module is used by each nucleus. The usage of each peak module in single nuclei was examined, and peak modules exhibiting high usage only in one batch were ignored (**Figure 2B** and **Supplementary Figure 2B**). Eventually, five peak modules (peak modules 2, 3, 4, 6 and 10) with consistent and reproducible usage across batches were used for the downstream analysis (**Supplementary Table 2**).

**Figure 2.**
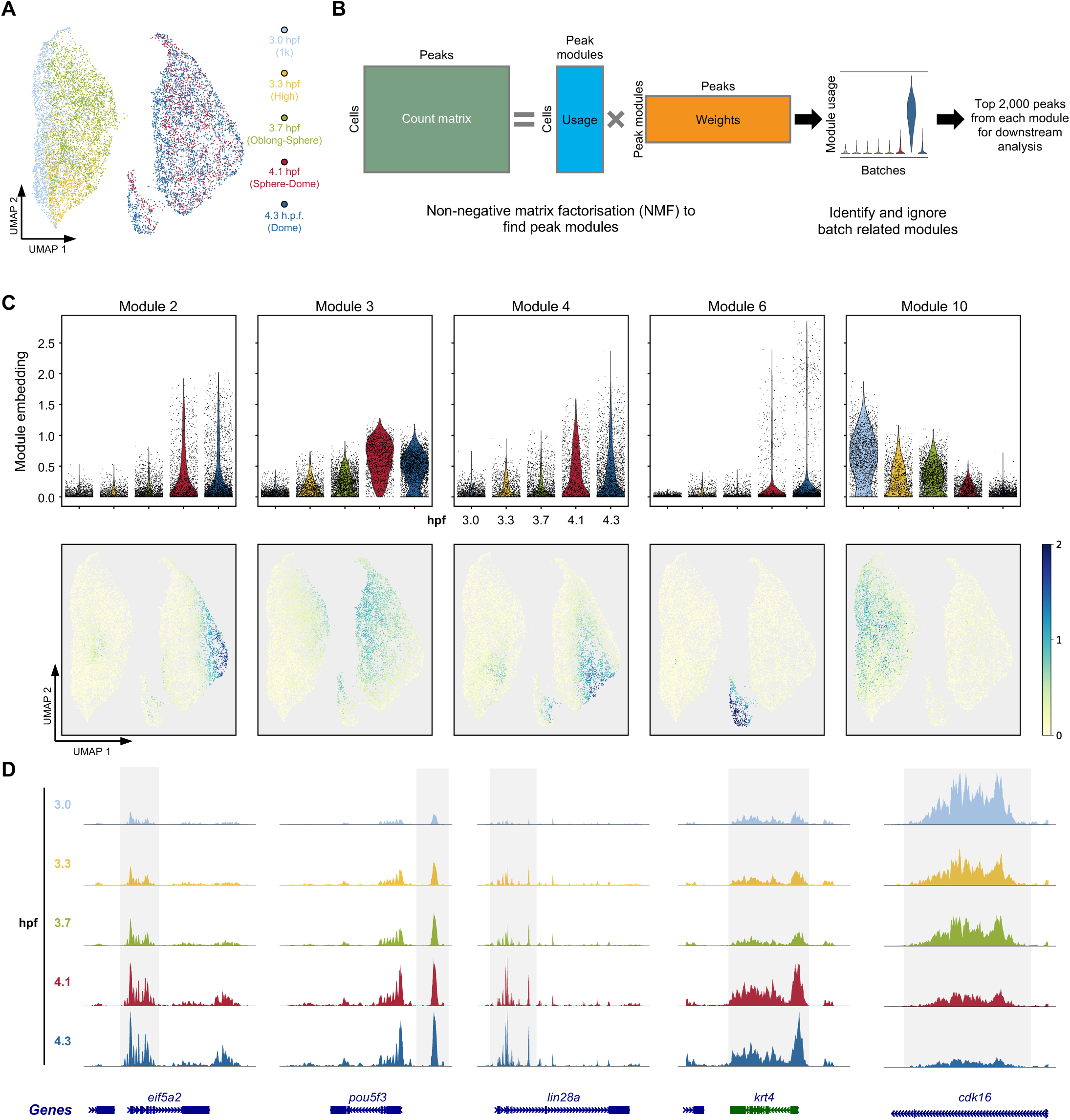
The dynamics of open chromatin peaks during early development. **(A)** The BBKNN integrated UMAP of all single-nucleus ATAC-seq profiles coloured by time points. **(B)** Schematic view of the non-negative matrix factorisation (NMF) workflow to identify peak modules that covary across nucleus in the whole data. Only reproducible peak modules were used for the downstream analysis. Batch-related peak modules were ignored. **(C)** The embedding values of peak modules 2, 3, 4, 6 and 10 in each single nucleus shown as the violin plot (top) and on UMAP (bottom). **(D)** Browser tracks of the single-nucleus aggregate of ATAC-seq signals from typical examples of each peak module in **(C)**.

Consistent with the notion that chromatin accessibility is gradually established during ZGA in zebrafish (12–14) and other species (42–47), the overall scATAC-seq enrichment around the transcriptional start sites (TSS) was increasing throughout the time course (**Supplementary Figure 2C**). The usages of most peak modules (modules 2, 3, 4, and 6) were increasing during the developmental time course and reached maximum in different subsets of cells at later stages (**Figure 2C**). The examination of the aggregates of scATAC-seq reads showed peaks within these modules became gradually accessible during the development process (**Figure 2D**). Interestingly, a peak module (module 10) with decreasing usage was also identified (**Figure 2C**), indicating certain genomic regions are turning into an inaccessible state following

ZGA (**Figure 2D**).

To further investigate the characteristics of the five peak modules, public ChIP-seq experiments of important histone marks and histone variants across different developmental stages in zebrafish were collected (48–50). A hidden Markov model, ChromHMM (51), was used to categorise the zebrafish genome into eight different states based on the ChIP-seq data, including various types of promoters and enhancers (**Figure 3A**). The distributions of the five peak modules across the chromatin states were quite different. Modules 2 and 3 were exclusively located in the constantly active promoter regions (**Figure 3A**, ChromHMM state 1), and modules 4 and 6 were mainly located in the enhancer regions (**Figure 3A**, ChromHMM states 3, 5 and 6).

**Figure 3.**
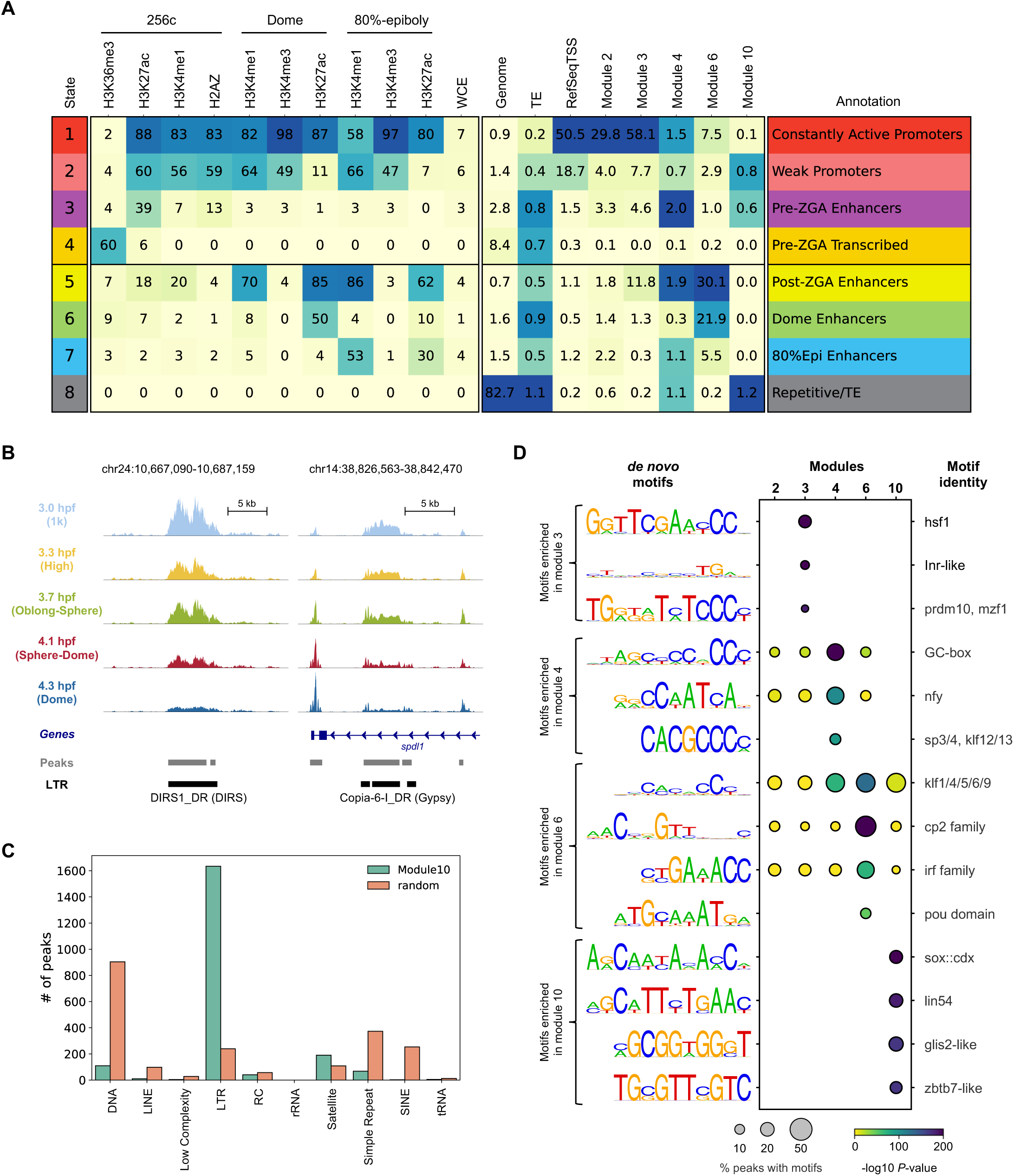
Distinct genomic features are associated with peaks from different peak modules. **(A)** ChromHMM analysis of the zebrafish genome using previously published ChIP-seq data sets. Peaks from peak modules 2, 3, 4, 6 and 10 were also compared to the chromatin states defined by the combinations of histone marks or variants. **(B)** Genome browser tracks showing two examples for peaks from peak module 10. **(C)** The distribution of the number of peaks in peak module 10 that overlapped the indicated transposable element class. Random was the background, which contained 2, 000 randomly selected peaks from all ATAC-seq peaks. **(D)** HOMER *de novo* motif results from each peak module. The percentage of peaks containing the motif and the p-values were shown. The best match to known transcription factor motifs was shown at the right-hand side.

Consistently, peaks from modules 2 and 3 were close to the TSS of annotated genes, and most peaks from modules 4, 6 and 10 were located in the intergenic regions (**Supplementary Figure 3A**). Notably, module 6 was preferentially located in the dome-specific enhancers (ChromHMM state 6) and the post-ZGA stage enhancers (ChromHMM state 5). Considering the usage of module 6 was most prominent in a subset of cells during the late stages (**Figure 2C**), peaks within module 6 were likely to play an important role in cell type specification (discussed later). Gene ontologies using GREAT (52) also suggested distinct functions for genes associated with each peak module. For example, the most enriched biological processes in modules 2 and 3 are translation and chromatin assembly (**Supplementary Figure 3B**), indicating these processes are critical throughout early embryo development which is consistent with the general knowledge (1).

One interesting observation was that module 10 was mainly located in the regions where transposable elements (TEs) reside (**Figure 3A**, ChromHMM state 8). Visual inspections on individual TEs showed the scATAC-seq reads covered the entire TE, and the chromatin status around the TEs became gradually closed after ZGA (**Figure 3B**). A more detailed analysis on TE classes and families showed >80% of the top 2, 000 peaks from module 10 overlapped long terminal repeats (LTR) (**Figure 3C**). Within all LTRs from module 10, the vast majority were dictyostelium intermediate repeat sequence (DIRS) (**Supplementary Figure 3C**). Intriguingly, a recent TE analysis (53) of gene expression data (54) during zebrafish early embryogenesis indicated LTRs were highly expressed immediately following ZGA, but the dynamic of the expression was not concordant with the chromatin accessibility changes (**Figure 2C** and **Supplementary Figure 3D**). These results together suggest a more complex regulatory mechanism of LTRs from module 10.

*De novo* motif analysis (31) revealed different transcription factor motifs enriched within the five peak modules (**Figure 3D**). While no prominent motifs were identified in module 2, motifs bound by factors including Hsf1 (55) and Nfya (44) that were important for ZGA reported in mice were more enriched in modules 3 and 4, respectively (**Figure 3D**). In addition, several motifs were identified as specifically enriched in peaks from module 10. When the DNA methylation profiles were examined around these LTRs within module 10 at 3 hpf (1k stage) from a recent study (56), they were quite different from DNA methylation profiles of typical open chromatin regions (**Supplementary Figure 3E**). Therefore, the exact meaning and functional implications of reduced chromatin accessibility of LTRs in module 10 during development need further investigation.

### Distinct chromatin signatures of cell types emerge around the sphere and dome stages

Since discrete cell populations started to show up around the 4.1 hpf (sphere-dome) stage (**Figure 1C**) and cells from the 4.1 hpf and 4.3 hpf (dome) had similar chromatin profiles (**Figure 2A**), all nuclei (a total of 4, 376 nuclei) from these two late stages were taken out for a more detailed analysis in order to identify emerging cell types. Clustering on the data from the 4.1 hpf and the 4.3 hpf time points revealed four different cell clusters (**Figure 4A** and **Supplementary Figure 4A**). Based on the genes around the differentially accessible peaks in each cluster, two of the clusters were identified as the enveloping layer (EVL) cells, a type of differentiated cells that covers the outside of the embryo (57), and the yolk syncytial layer (YSL) cells which are an extra-embryonic tissue that contributes to the patterning of the embryo (58) (**Figure 4A**). High chromatin accessibility around canonical marker genes, such as *krt18a.1*, *krt4*, *cldne* and *gata3*, were observed specifically within the EVL cell cluster (**Figure 4B**, **Figure 4C** and **Supplementary Figure 4B**). Consistent with previous studies that *afp4* and *cyp11a1* were specifically expressed in the YSL cells during early embryogenesis (59, 60), clear open chromatin peaks around the *afp4* and *cyp11a1* loci were observed only in the YSL cell cluster (**Figure 4B** and **C**). The rest of the two cell clusters (**Figure 4A,** DEL1 and DEL2)constituted the majority of the cells and were most likely to be the undifferentiated pluripotent cells in the deep layer (DEL) between EVL and YSL. Although both were considered to be located in the deep layer, differences between DEL1 and DEL2 were clearly observed. Many peaks, especially those from module 2, showed higher accessibility in cells from the DEL2 cluster (**Supplementary Figure 4A**). These peaks were mainly associated with genes involved in translation, mitotic cell cycle, mRNA metabolic process and RNA splicing (**Supplementary Figure 3B**), suggesting DEL2 cells may have higher activity in those biological processes compared to cells in the DEL1 cluster.

**Figure 4.**
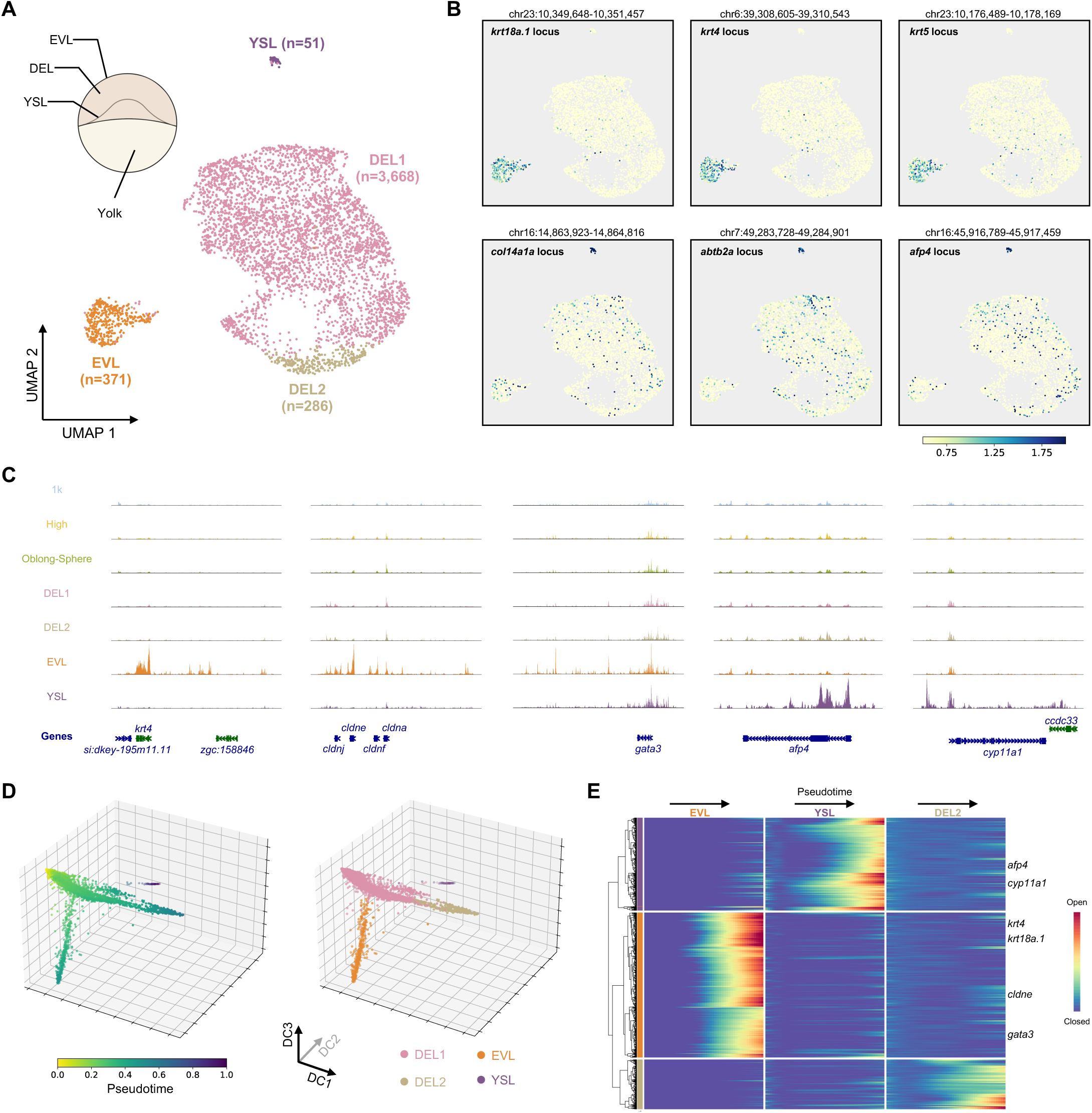
Emerging different chromatin profiles among single cells when entering the dome stage. **(A)** UMAP visualisation of nucleus from the 4.1 hpf (sphere-dome) and the 4.3 hpf (dome) stages coloured by different clusters. Cluster annotations were indicated on the UMAP plot. A schematic view of the structure of the embryo was also shown at the top left. **(B)** The same UMAP plot as in **(A)**, coloured by the normalised reads from the indicated gene locus. **(C)** Genome browser tracks showing the aggregated single-nucleus ATAC-seq signal from indicated cell clusters. Five examples were shown around the mark genes of EVL and YSL cells. **(D)** Diffusion pseudotime analysis of the nuclei from the 4.1 hpf (sphere-dome) and the 4.3 hpf (dome) stages, coloured by the pseudotime (left) or cell identity (right). **(E)** Heatmap representation of gene activity changes along three different branches. Important marker genes for EVL and YSL cells were indicated.

We also examined the expressions of the key marker genes around similar stages from a scRNA-seq atlas of zebrafish development (4). The expression dynamics of EVL marker genes were concordant with the chromatin accessibility changes during the time course, where those genes remained silent before the dome stage and only started to express in the EVL subset of cells in the dome stage (**Supplementary Figure 4C**). Since the YSL cells were not identified in the scRNA-seq atlas (4), some marker genes such as *afp4* were not detected. The YSL marker gene *cyp11a1* was detected as early as the high stage, possibly due to the fact it is a maternal transcript (60). The discrepancy of the YSL cells between scRNA-seq and scATAC-seq data was probably due to technological difference (see **Discussion**).

To further explore the relationship among those clusters during development, nuclei from the same cluster were first pooled and peak calling was performed for each cluster aggregate (see **Methods**). Then the resulting cell-by-peak matrix were used for the trajectory inference with diffusion pseudotime (DPT) (36). The DPT analysis split the cells into three different trajectories (**Figure 4D**). The main trajectory from DEL1 to DEL2 represented the differentiation of pluripotent cells which ultimately would lead to the formation of all other body parts of the zebrafish. Two branches emerged from the main trajectory represented the EVL and YSL fate acquisition, respectively (**Figure 4D**). The pseudotemporal ordering of cells also provided key dynamic information of open chromatin sites along each branch (**Figure 4E**). For example, in the EVL branch, the regulatory regions around key marker genes *krt4*, *krt18a.1* and *gata3* became fully open at the end of the pseudotime, but the *krt4* and *krt18a.1* loci started to gain accessibility earlier than *gata3* (**Figure 4E**).

### Newly identified cell-type specific enhancers and transcription factors drive the EVL and YSL cell fate acquisitions

We noticed the peak modules 2 and 6 identified by NMF reached the highest usage in the DEL2 and EVL cell clusters, respectively (**Supplementary Figure 4A**). In order to systematically find marker peaks of each cell cluster, differential accessibility analysis was performed among peaks called by the cell cluster aggregate in the previous section (**Supplementary Table 3**). While very few specific peaks were found for the DEL1 or DEL2 cell cluster, many novel marker peaks that were accessible only in the EVL or YSL cell cluster were identified (**Supplementary Table 3**). Compared to the eight-state ChromHMM model (**Figure 3A**), both the EVL and the YSL marker peaks were mainly located in the dome specific enhancer and the post-ZGA stage enhancer regions (**Figure 5A**). A more detailed investigation of ChIP-seq reads of key histone modifications (50) around the marker peaks indicated low level of H3K4me3 and high levels of H3K4me1 and H3K27ac around the centre of EVL and YSL marker peaks (**Figure 5B**). The H3K4me1 and H3K27ac signals around the marker peaks were observed at the dome stage and persisted in the 80% epiboly stage, indicating these newly identified marker peaks by our scATAC-seq data are putative enhancers that are important for the cell fate specification at the dome stage and remain active at least through the 80% epiboly stages.

**Figure 5.**
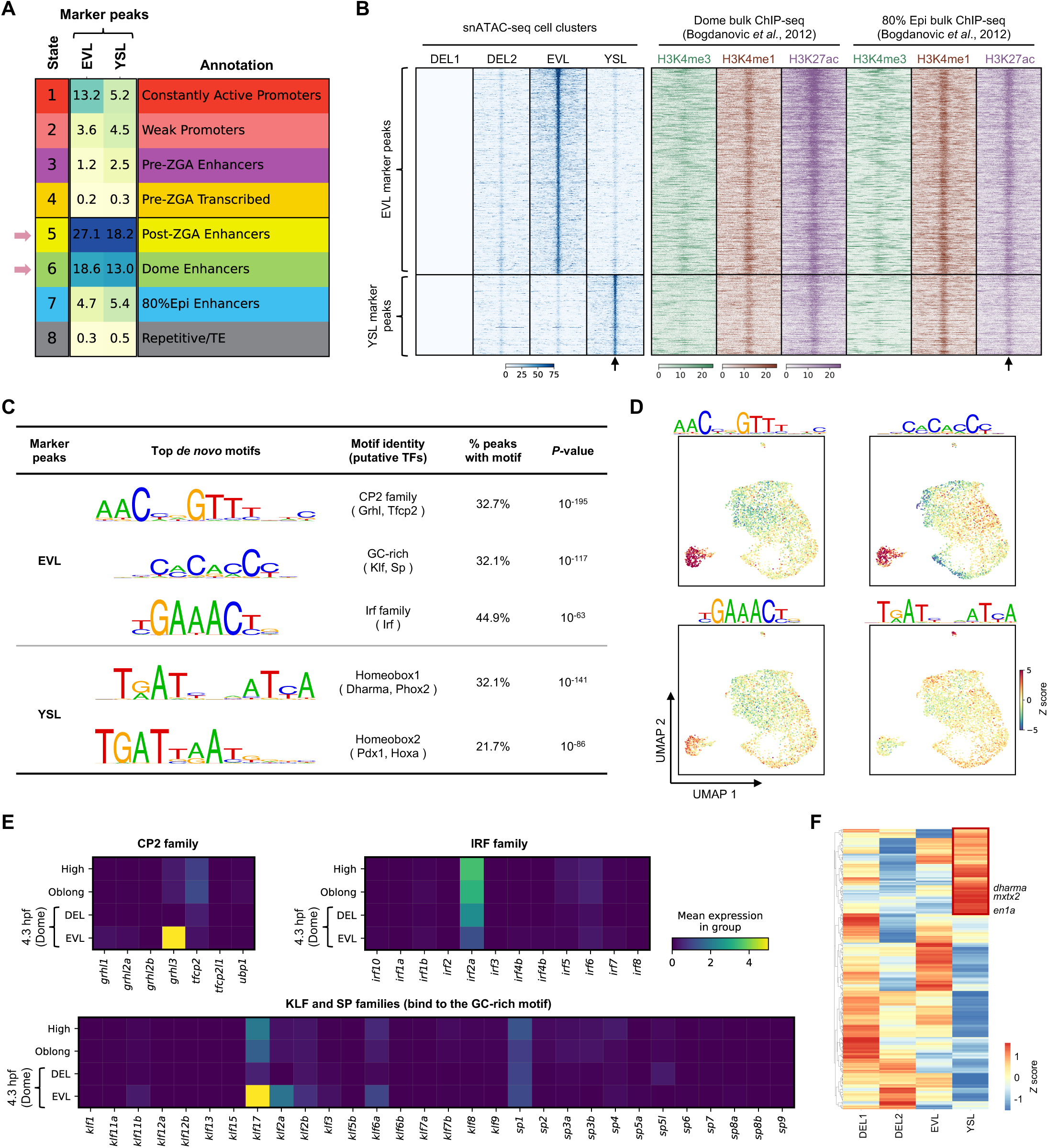
Putative enhancers located in the marker peaks of EVL and YSL cells. **(A)** The comparison of the EVL and YSL marker peaks to the eight chromatin states described in Figure 3A. **(B)** Heatmap representation of the aggregated of scATAC-seq reads (left) or public ChIP-seq reads of indicated histone marks (right) around EVL and YSL marker peaks. Black arrows indicate the centre of the peak. 5 kb upstream and downstream of the peak were plotted. **(C)** HOMER *de novo* motif discovery results of the marker peaks of EVL and YSL cells. **(D)** The same UMAP plot as in Figure 4A, coloured by the deviation scores of indicated motifs calculated by chromVAR. **(E)** The mean expression levels of indicated genes in each stage from a previously published scRNA-seq atlas. All members from the CP2, the IRF, the KLF and the SP family were investigated. **(F)** Heatmap representation showing the gene activity scores of all homeobox transcription factors measured by scATAC-seq data in different cell clusters. Red box indicated the 73 homeodomain factors that have higher gene activity scores in the YSL cluster.

To find out potential transcription factors that shape the chromatin profiles of the EVL and YSL cells, *de novo* motif discovery was performed on the two sets of marker peaks, respectively. The most enriched motif from the EVL marker peaks resembled the DNA binding consensus of the CP2 family, which could be bound by transcription factors such as Grhl and Tfcp2 (**Figure 5C**). A GC-rich motif that was similar to the Klf family and Sp (specificity protein) family consensus and a third one that resembled the IRF family motif were also found enriched within the EVL marker peaks (**Figure 5C**). Motif activity inferred by chromVAR (33) confirmed the results that the aforementioned motifs had the highest activity in the EVL cells (**Figure 5D**). The motif results point to the possibility that the CP2, Klf, Sp and Irf families might play an important role in shaping the chromatin state of EVL cells.

Two versions of the homeobox motifs were found over-represented within the YSL marker peaks (**Figure 5C** and **D**). The top motif, homeobox 1, was palindromic and consisted of the half-site TRATY with a 1-bp spacer, where R stands for A or G and Y stands for C or T (**Figure 5C**). The motif was previously shown to be bound by the mouse homeodomain transcription factors, such as Cphx1 and Phox2a/b (61–63), though no orthologs of the *Cphx1* gene was present in zebrafish. Another version of the homeodomain motif, homeobox 2, contained the non-canonical core sequence TGAT followed by the canonical homeobox core TAAT (**Figure 5C**) (61, 64). The homeobox 2 motif was shown to be bound by transcription factors such as Pdx1 and Hoxa (61, 64–66). These results suggested the homeodomain transcription factors were important to establish the chromatin signature of the YSL cells. Interestingly, *dharma*, a homeodomain transcription factor expressed in the YSL has previously been shown to be important for the function of YSL cells (58, 67), and it can potentially bind to the half-site of the homeobox 1 motif (66).

In order to narrow down the key transcription factors that are important for the establishment of the EVL and YSL fates, the expression levels of all members from relevant transcription factor families (68) were checked using the scRNA-seq atlas of early zebrafish embryo development (4). Since YSL cells were not present in the scRNA-seq atlas, we first focused on the gene expression of relevant transcription factor families in EVL cells. Intuitively, a transcription factor must be expressed in the cell to be functional. Therefore, we checked expression levels of all transcription factors that can potentially bind to the three main motifs discovered from the EVL mark peaks (**Figure 5E**). Within the CP2 family, *grhl3* is the only member that could be detected, and it was specifically expressed in EVL cells at the dome stage (**Figure 5E**). Similarly, *klf17* had the highest expression level in EVL cells at the dome stage, though *klf2*, *klf6a* and *sp1* that are able to bind to similar DNA motif could also be detected at the dome stage (**Figure 5E**). In the Irf family, only the mRNA of *irf2a* could be detected, but its expression level went down since the high stage (**Figure 5E**). Previous functional studies implicated the importance of Grhl3 and Klf17 during the development of EVL cells (69, 70). Combined with our scATAC-seq data, the results indicated that those transcription factors promote EVL development and functions through the establishment of EVL specific chromatin profiles at enhancer regions. Next we systematically investigated the gene activity scores of all homeodomain transcription factors in the YSL cluster by measuring their promoter plus gene body accessibilities (35). A total of 73 homeodomain transcription factors were found to have much higher gene activity scores in the YSL cells compared to other cell clusters (**Figure 5F**, red box and **Supplementary Table 4**), including *dharma* and *mxtx2* that were previously shown to be important for the development of YSL (67, 71). Interestingly, *en1a*, a predicted paralog of mouse *Cphx1*, was also found to have higher gene activity score in the YSL cells (**Figure 5F**). This analysis suggests the 73 homeodomain transcription factors might be potentially expressed in the YSL cells and could possibly bind to the homeobox motifs enriched within the YSL marker peaks.

Finally, co-accessibility analysis (35) was performed to link distal regulatory elements to their potential target genes. In this way, many putative distal enhancers were linked to important marker genes in EVL and YSL cells, respectively (**Figure 6** and **Supplementary Figure 5**). Many of the enhancers have at least one previously described motif and reproducible H3K27ac signals from two independent studies (49, 50) (**Figure 6**). This demonstrated that our data have identified many potential cell-type specific enhancers that may play critical roles in the cell fate specifications of EVL or YSL cells.

**Figure 6.**
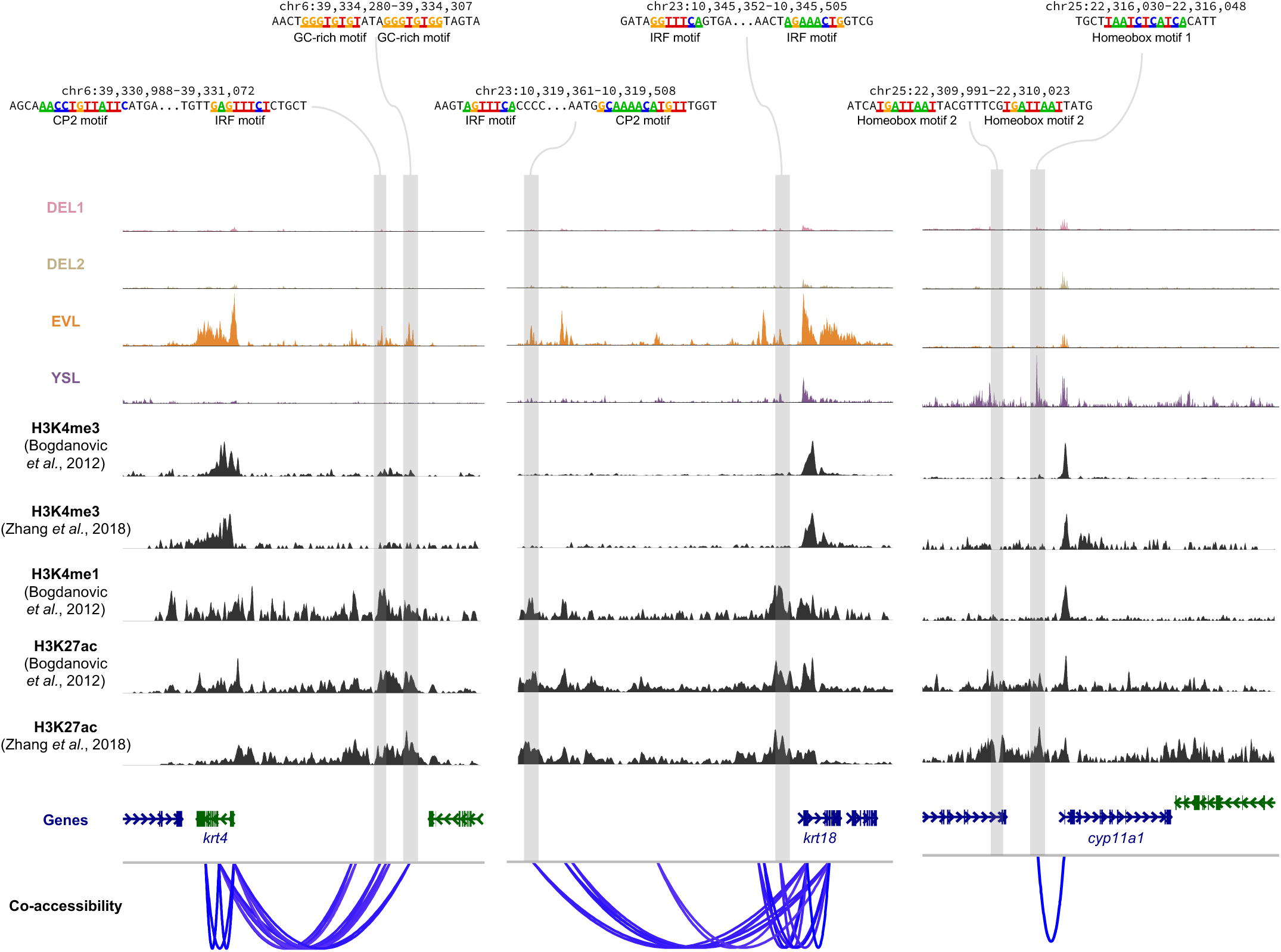
Co-accessibility analysis links putative enhancers to their target genes. Examples of co-accessibility of peaks around EVL and YSL marker genes. The linkage strength analysed by Cicero was shown at the bottom. Cell-type specific open chromatin peaks with motifs from Figure 5C were highlighted. ChIP-seq data of H3K4me3, H3K4me3 and H3K27ac from two previous studies were also shown.

## Discussion

The acquisition of cell-type specific molecular signatures during early development is a key feature in all multi-cellular organisms. Recent technological development has allowed the interrogation of various molecular profiles at the single cell level, including gene expression, chromatin accessibility and chromosome conformation etc (72, 73). The techniques offer excellent opportunities to study cell fate specifications during embryo development. In zebrafish, several single-cell atlases have been generated in terms of gene expression (4–10) and chromatin accessibility (15–17). They mainly focused on either the late stages of embryogenesis (mostly after gastrula) or adult tissues, partly due to the fact that early embryo development is driven by maternal factors in zebrafish (2).

In this study, we investigated the chromatin accessibility landscapes at the single-nucleus level, with over 10, 000 nuclei across five key stages that span the first hour immediately following ZGA. We identified important open chromatin peak modules with distinct sequence features during the developmental time course (**Figures 2** and **3**). More importantly, the data suggested cells started to gain distinct chromatin signatures when they progressed from the sphere to the dome stage. Cell-type specific distal regulatory elements for EVL and YSL cells, respectively, were identified using our scATAC-seq data (**Figures 4** and **5**). They were marked by low-level of H3K4me3 and high level of H3K4me1 and H3K27ac and contained important transcription factor binding motifs (**Figures 5** and **6**), indicating those distal regulatory elements are putative enhancers that are important for the EVL and YSL cell fate specifications.

The YSL cells were not present around the dome stage in the previous two scRNA-seq atlases (4, 5). Since the YSL cells are present as nuclei in the syncytium on the surface of the yolk cell (58), it is likely that the typical single-cell isolation procedures in the scRNA-seq experiment would result in the loss of those nuclei during the experiments. The ChIP-seq data of histone marks used to annotate the cell-type specific regulatory regions were taken from bulk experiments at the dome stage (49, 50). Our single-cell data suggested the ChIP signals of those histone marks might come from only a subset of cells, EVL or YSL, at the dome stage, and the bulk ChIP-seq experiments seem to be sensitive enough to enrich and uncover those signals (**Figure 5B** and **Figure 6**). With the development of single-cell ChIP-like techniques (74–83), it is possible to characterise potential cell-type specific enhancers more accurately in the future.

Our scATAC-seq data showed that EVL and YSL cells emerged and acquired distinct chromatin profiles at 4.1 hpf (**Figure 1C** and **Figure 4A**), indicating those cells start to differentiate and take specific cell fates. The classic cell fate commitment study from Ho and Kimmel showed cells at the dome stage remain pluripotent and uncommitted (84). When examined the chromatin status of key pluripotent genes, including *pou5f3*, *nanog*, *sox19b*, in different cell clusters at the dome stage, all of them exhibit a transcriptional permissive state with many open chromatin peaks covering the gene loci (**Supplementary Figure 6**). These observations suggest that even EVL and YSL cells start to obtain specific chromatin features that are distinct from DEL cells, their cell fates are not fully committed yet at the dome stage.

In general, our data serve as a useful resource that provides an overview of the *cis*-regulatory dynamics during the first cell fate specification of zebrafish early embryogenesis. Combined with functional studies, gene expression profiles and other data, it allows more detailed investigation of the mechanism underlying cell fate decisions.

## Data availability

The raw sequencing data, the processed peak by cell count matrix, the R object and the meta data have all been deposited at ArrayExpress under the accession number E-MTAB-13115.

## Acknowledgement

We thank all members from the Chen and Jin labs for the helpful discussion of the project. We also thank Dr. Zilong Wen for the helpful suggestions on the writing of the manuscript. We thank Xibin Lu for the excellent support of FACS. We acknowledge the assistance of SUSTech Core Research Facilities. The computational work was supported by Center for Computational Science and Engineering at Southern University of Science and Technology. Part of Figure 1a and Figure 4a were created with BioRender.com.

## Author contributions

X.C and N.H conceived the project. Q.X, Y.Z, W.X, N.H and X.C designed the experiments. Q.X performed the experiments with the help from W.X and N.H. Y.Z, carried out the computational analysis with the help from X.C, Q.X and N.H. D.L, J.W., X.C and N.H supervised the entire project. All authors contributed to the writing.

## Code availability

The code used for the data processing and analysis mentioned in the method section is available on the GitHub repository https://github.com/helianfeixing/Zebrafish_scATAC

---

**Supplementary Figure 1.**
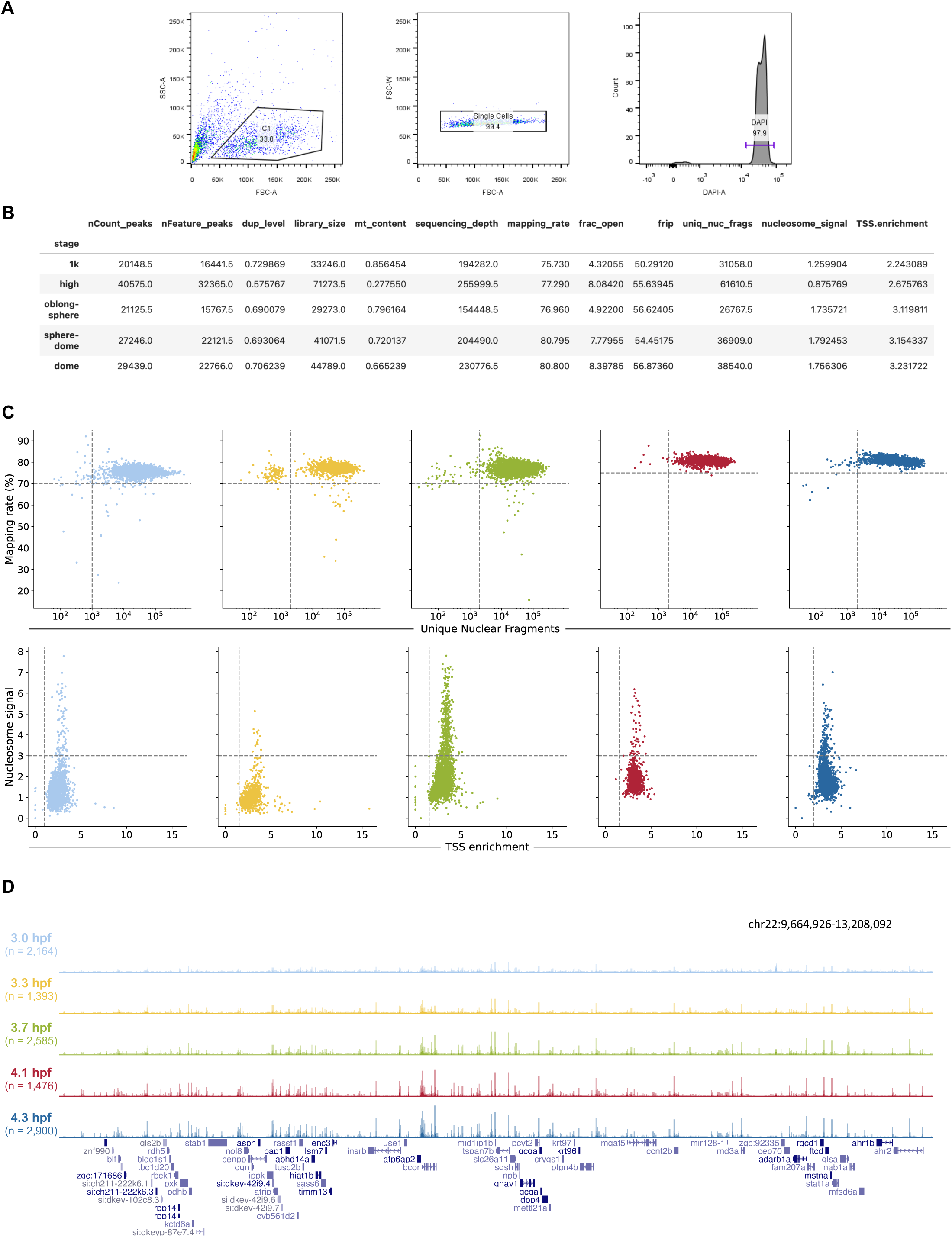
Quality control (QC) of the whole single-nucleus ATAC-seq experiment. **(A)** Examples of FACS for a typical experiment. FSC-A and SSC-A were used to remove cell debris. FSC-A and FSC-W were used to select only single nucleus. DAPI+ events were used to select only successfully permeabilised cells. **(B)** Median information of each QC metric in different stages. **(C)** QC cutoff used to remove failed wells in the experiment. **(D)** UCSC genome browser tracks from the indicated genomic locus showing an overview of the data quality.

**Supplementary Figure 2.**
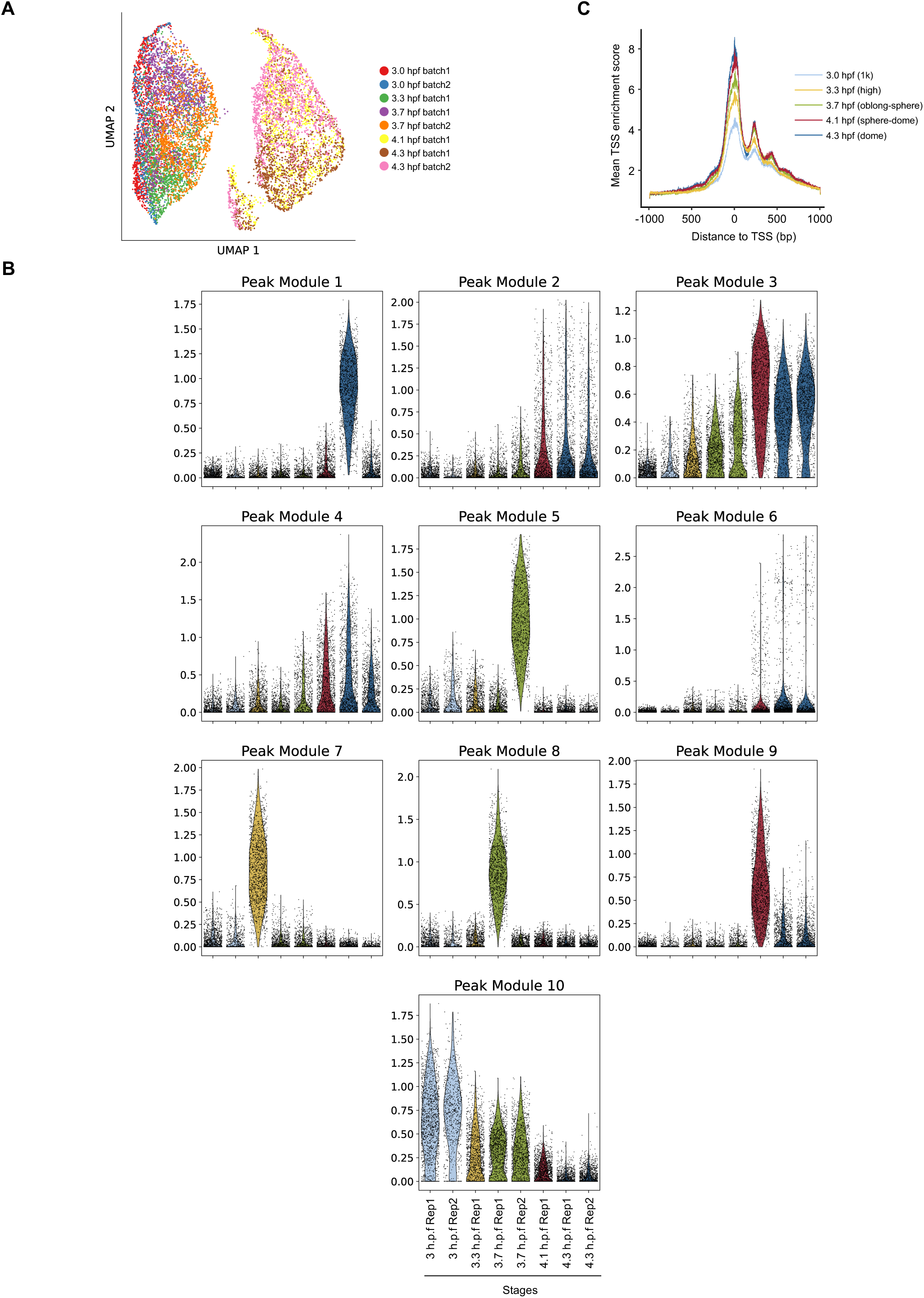
The overview of the entire single nucleus ATAC-seq data set. **(A)** BBKNN integrated UMAP of all nucleus, coloured by experimental batches. **(B)** The module weights of each peak module grouped by experimental batches. **(C)** The global TSS enrichment scores of each time point.

**Supplementary Figure 3.**
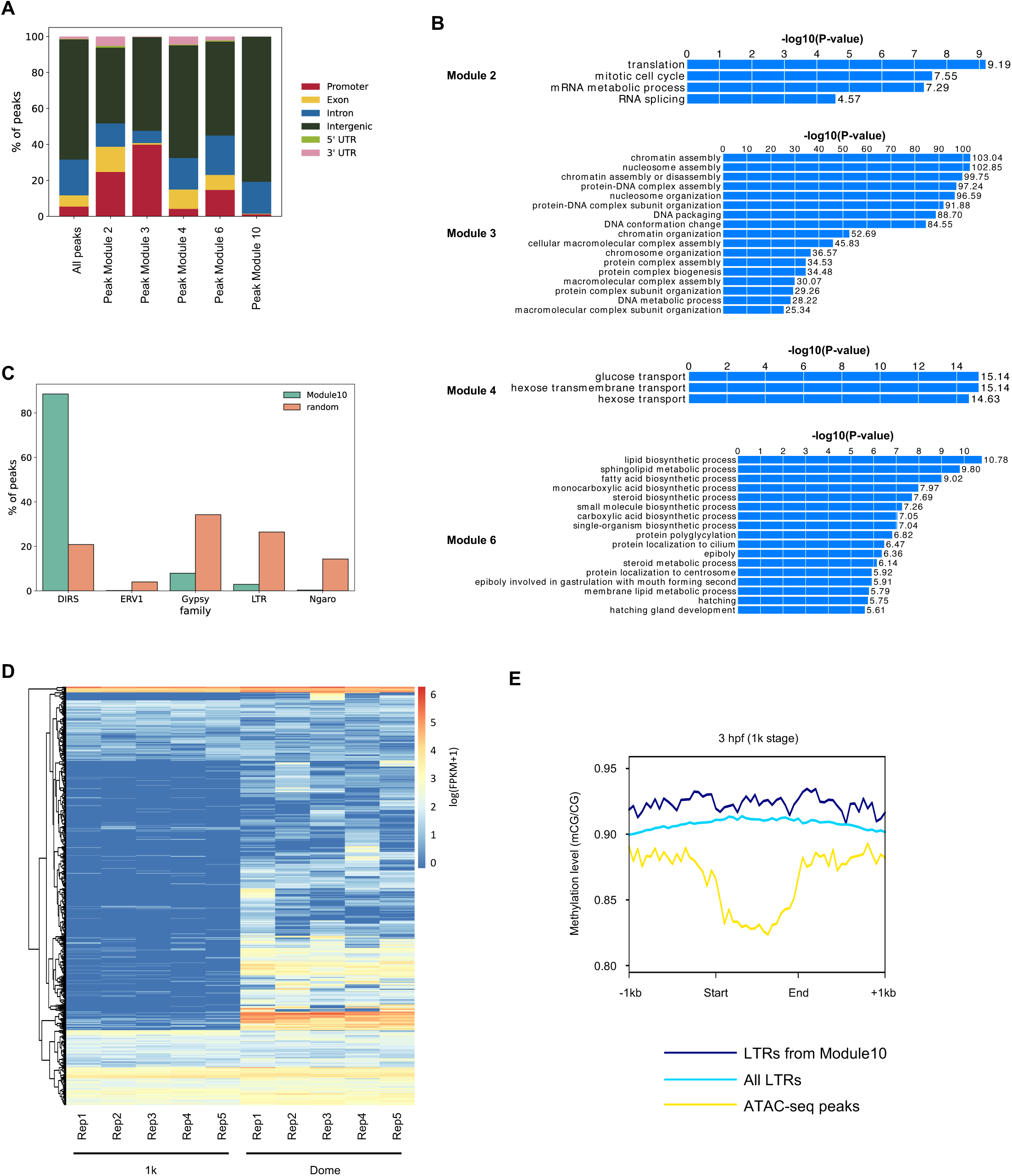
Different genomic features and functions of each peak module. **(A)** The genomic distribution relative to annotated genes of each peak module. **(B)** GREAT analysis of enriched biological processes in each peak module. **(C)** The distribution of the number of peaks in peak module 10 that overlapped the indicated LTR family. Random was the background, which contained 2, 000 randomly selected peaks from all ATAC-seq peaks. **(D)** The expressions of 519 LTRs in module 10 at 1k and dome stages. Data were taken from White *et al.* 2017 (54). **(E)** The DNA methylation levels (mCG/CG) at 3 hpf (1k stage) around indicated regions. 1 kb upstream and downstream were also plotted. Data were taken from Ross *et al.* 2023 (56).

**Supplementary Figure 4.**
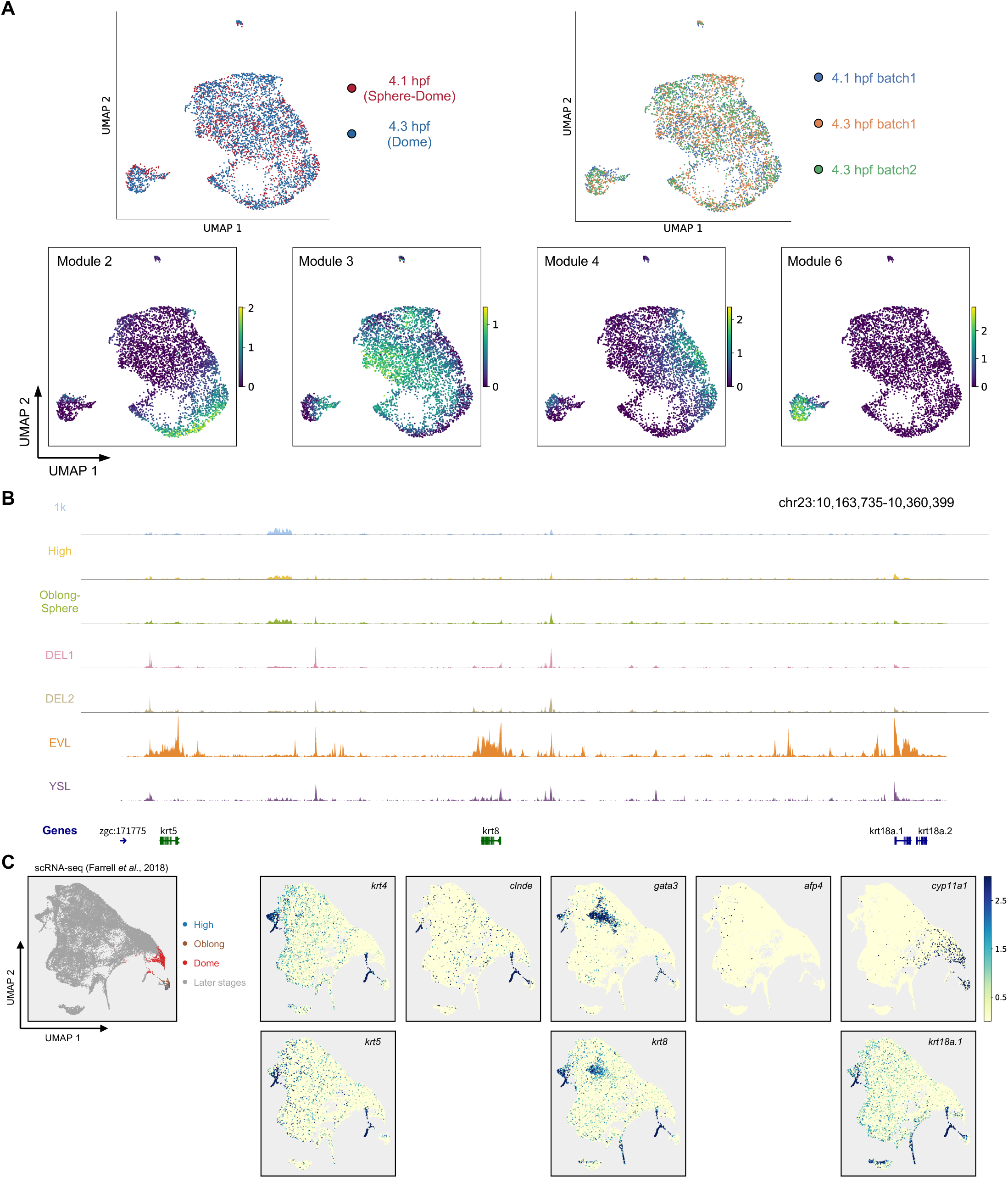
Analysis of nucleus from 4.1 hpf (sphere-dome) and 4.3 hpf (dome) stages. **(A)** The same UMAP plot as in Figure 4A, coloured by time point (top left), experimental batches (top right) and module weights (bottom). **(B)** Browser tracks showing specific peaks around three EVL marker genes. **(C)** The expression levels of indicated genes from a previously published scRNA-seq atlas.

**Supplementary Figure 5.**
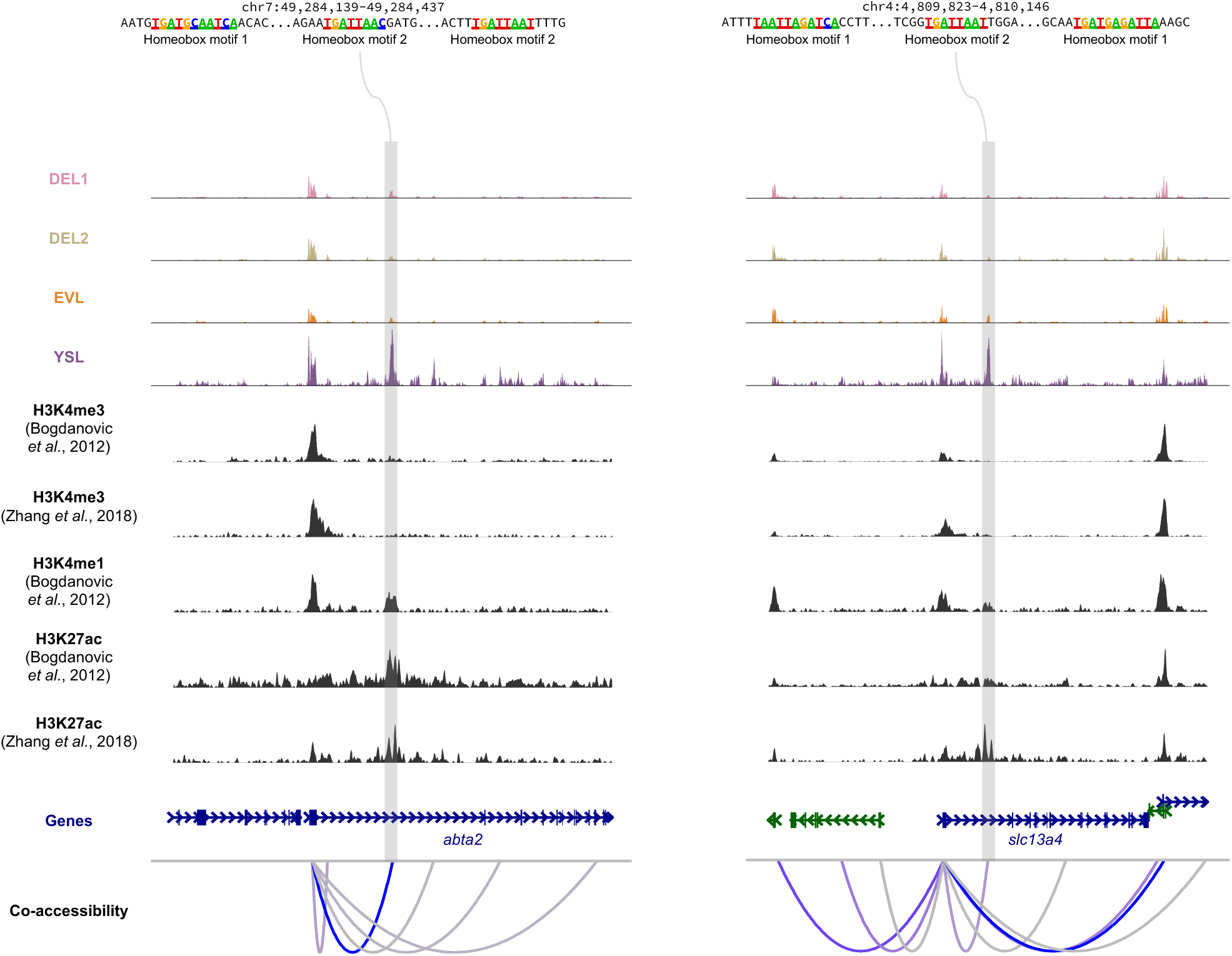
Examples of peak co-accessibility around two YSL marker genes. Cell-type specific open chromatin peaks with motifs from Figure 5C were highlighted. ChIP-seq data of H3K4me3, H3K4me3 and H3K27ac from two previous studies were also shown.

**Supplementary Figure 6.**
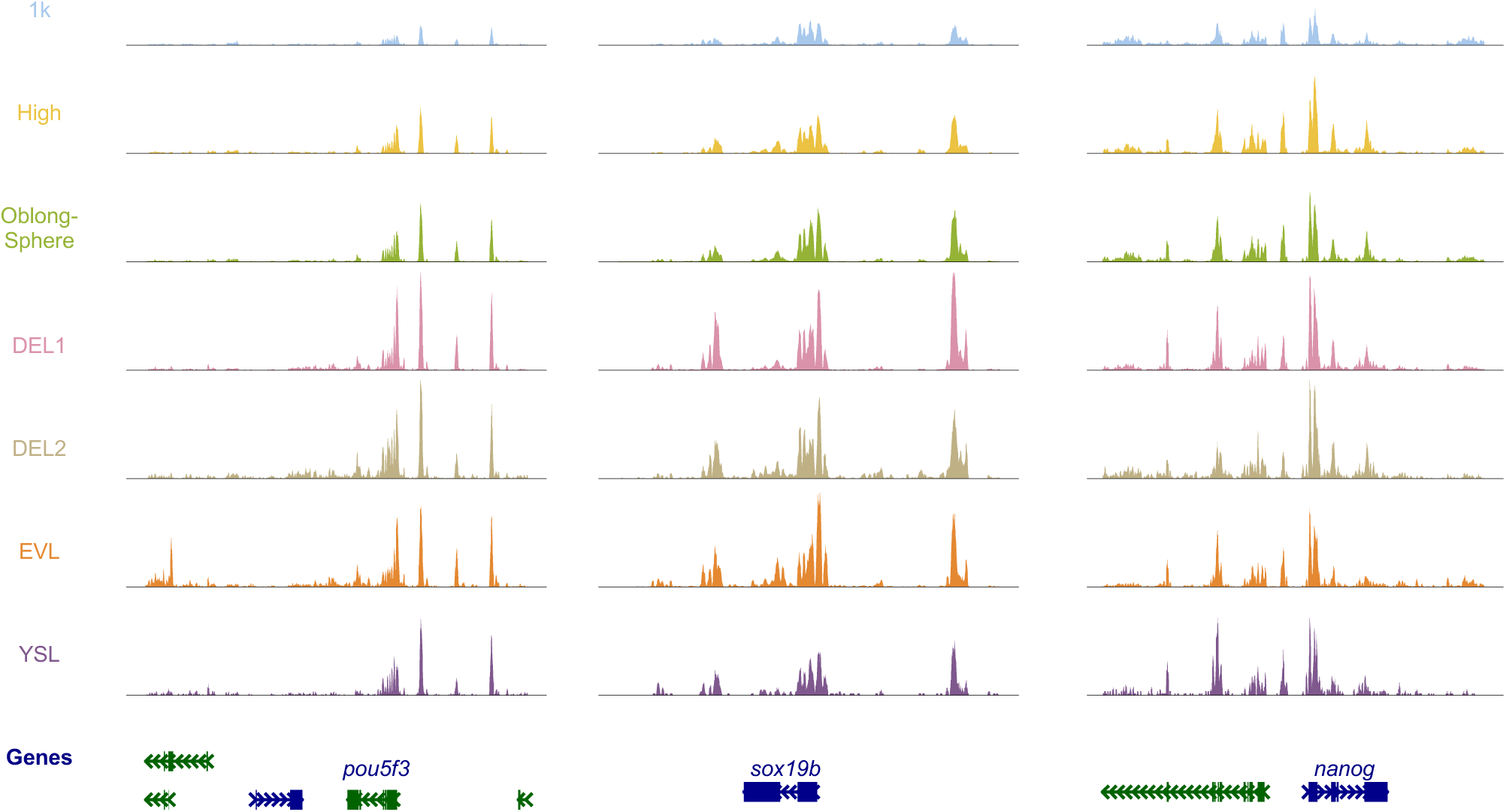
Browser tracks showing the aggregated single-nucleus ATAC-seq signal in each cell cluster around the three pluripotent genes.

